# Muscle cofilin alters neuromuscular junction postsynaptic development to strengthen functional neurotransmission

**DOI:** 10.1101/2023.11.21.568166

**Authors:** Briana Christophers, Shannon N. Leahy, David B. Soffar, Victoria E. von Saucken, Kendal Broadie, Mary K. Baylies

## Abstract

Cofilin, an actin severing protein, plays critical roles in muscle sarcomere addition and maintenance. Our previous work has shown *Drosophila* cofilin (*DmCFL*) knockdown causes progressive deterioration of muscle structure and function and produces features seen in nemaline myopathy (NM) caused by cofilin mutations. We hypothesized that disruption of actin cytoskeleton dynamics by *DmCFL* knockdown would impact other aspects of muscle development, and, thus, conducted an RNA sequencing analysis which unexpectedly revealed upregulated expression of numerous neuromuscular junction (NMJ) genes. We found that DmCFL is enriched in the muscle postsynaptic compartment and that DmCFL deficiency causes F-actin disorganization in this subcellular domain prior to the sarcomere defects observed later in development. Despite NMJ gene expression changes, we found no significant changes in gross presynaptic Bruchpilot active zones or total postsynaptic glutamate receptor levels. However, *DmCFL* knockdown results in mislocalization of glutamate receptors containing the GluRIIA subunit in more deteriorated muscles and neurotransmission strength is strongly impaired. These findings expand our understanding of cofilin’s roles in muscle to include NMJ structural development and suggest that NMJ defects may contribute to NM pathophysiology.

**Summary statement:** Cofilin regulates muscle postsynaptic actin organization, structural maintenance, glutamate receptor composition, and neuromuscular junction function in a *Drosophila* nemaline myopathy disease model.

## Introduction

Skeletal muscle is critical for daily functions, including respiration, posture control, and coordinated movement. Muscle contraction and relaxation require many levels of organization: each cell is made up of myofibrils, which are concatenations of the contractile sarcomeres. The sarcomere is composed of thin filaments anchored at Z-discs and thick filaments containing myosin motors that enable filament sliding. Thin filaments comprise filamentous actin (F-actin) and actin-binding proteins that anchor, maintain length, and regulate myosin binding.

The actin depolymerizing factor (ADF)/cofilin family maintains filament length by inducing a unique conformational change at ADP-actin filaments to promote actin severing and prevent G-actin nucleotide exchange (Galkin et al., 2011; Hayden et al., 1993). The vertebrate family includes ADF/destrin, cofilin-1, and cofilin-2, with cofilin-2 predominant in postnatal and mature skeletal muscle (Mohri et al., 2000; Ono et al., 1994; Vartiainen et al., 2002). Cofilin-2 is important for skeletal muscle development, maintenance, regeneration, and for determining thin filament length (Agrawal et al., 2012; Gurniak et al., 2014; Kremneva et al., 2014; Ono et al., 1994; Thirion et al., 2001). Mutations in twelve actin and sarcomere-related genes, including *CFL2,* have been implicated in nemaline myopathy (NM), a skeletal muscle disease that presents with progressive weakness, particularly during the perinatal period in its most severe forms (Christophers et al., 2022; Sewry et al., 2019). NM diagnosis is based on myopathic clinical disease hallmarks, including hypotonia and proximal muscle weakness, as well as the presence of actin-rich accumulations (known as nemaline rods or bodies) and myofibrillar disruption on pathology. *CFL2* NM patients have generalized muscle weakness; delayed motor skills; stiff spine; kyphoscoliosis; and joint contractures (Agrawal et al., 2007; Fattori et al., 2018; Ockeloen et al., 2012; Ong et al., 2014). Mutations in *CFL2* have been described as recessive and hypomorphs and lead to reduced levels of cofilin activity in muscle.

Cofilin is conserved across species, and cofilin-deficient disease models have been developed in mice, *C. elegans*, and *Drosophila melanogaster*. Several strategies have been used to study cofilin-2 in mouse models, including constitutive knockout, muscle-specific excision, chimeras that have a combination of knockout and wildtype cells, and knock-in of an NM mutation (Agrawal et al., 2012; Gurniak et al., 2014; Mohri et al., 2019; Morton et al., 2015; Rosen et al., 2020). Mice from all these models are small and have myofibrillar organization defects. Most die in the early postnatal period, which indicates that cofilin-2 is not essential for initial myofibrillogenesis but is important for postnatal muscle maintenance. In *C. elegans*, the cofilin isoform UNC-60B is required for incorporating actin into developing myofibrils, and mutants develop actin aggregates in body wall muscle (Ono et al., 1999; Ono et al., 2003). In *Drosophila*, we previously characterized the effect of muscle-specific knockdown (KD) of the cofilin homolog *twinstar*, the only cofilin homolog in *Drosophila* (Balakrishnan et al., 2020). *Drosophila* cofilin (*DmCFL*) KD leads to progressive muscle deterioration over the larval developmental stages. Actin accumulation occurs as a result of improper sarcomere addition at the growing muscle ends, and, eventually, they form throughout the cell, reminiscent of those seen in NM patients. Introducing wild type human *CFL1* or *CFL2* rescued this deterioration phenotype in *DmCFL* KD, while that of a human *CFL2* NM mutation did not, supporting the use of this system to understand human NM disease.

To further investigate the *DmCFL* KD model, we conducted RNA sequencing which identified changes in genes related to the neuromuscular junction (NMJ), the specialized synapse where the presynaptic motor neuron and postsynaptic muscle communicate. Early case reports examining NM muscle biopsies provide evidence of NMJ disruption, describing abnormal motor end plates and synaptic clefts (Fukuhara et al., 1978; Heffernan et al., 1968). Moreover, electromyography shows a myopathic pattern in NM patients, although it is unclear if there is additional superimposed NMJ transmission defect. Cytoplasmic actin and other actin-binding proteins are present at the vertebrate NMJ (Berthier and Blaineau, 1997; Bloch and Hall, 1983; Hall et al., 1981). Additionally, in vertebrates, actin podosomes are critical for shaping the complex organization of postsynaptic proteins (reviewed in Bernadzki et al., 2014). Thus, it is possible that disruption of actin and actin-binding proteins would affect the NMJ in NM patients. Recently, a case report of two NM patients with a mutation in alpha-actin describes abnormal postsynaptic muscle membrane at the NMJ, but it is unknown whether this finding is generalizable all NM patients (Labasse et al., 2022). At the widely-used *Drosophila* larval NMJ, proper actin organization at the muscle postsynapse is important for synaptic development, neurotransmitter receptor localization, and neurotransmission (Chen et al., 2005; Pielage et al., 2006; Wang et al., 2010). Based on the results of our RNA sequencing experiment and these previous *Drosophila* studies, we sought to better understand the NMJ in the *DmCFL* KD model and its contribution to disease development.

Here, we report that DmCFL is present near the muscle postsynaptic membrane, where it is important for proper actin organization in this region. We found that, in *DmCFL* KD, while the motor neuron properly innervates the muscle, postsynaptic structural organization is disrupted prior to and in parallel with muscle structure deterioration. Although transcriptomic analysis suggested changes in many synaptic components, we did not see defects in the presence of active zones or total glutamate receptors. However, DmCFL is important for the localization of glutamate receptor subunit A (GluRIIA) at the postsynapse, and *DmCFL* KD results in decreased NMJ neurotransmission. Together, these findings show that cofilin is important for NMJ postsynaptic structure, actin organization, GluRIIA localization, and synaptic signaling.

## Results

### Muscle-specific *DmCFL* knockdown alters neuromuscular junction (NMJ) gene expression

We have previously developed a *Drosophila* cofilin (*DmCFL)* knockdown (KD) model that has disrupted sarcomere structure, protein aggregates, and reduced muscle function, which are features of nemaline myopathy (NM) pathology (Balakrishnan et al., 2020). We used the GAL4/UAS system with a muscle-specific Myosin heavy chain promoter (*Mhc*-Gal4) driving an RNA interference (RNAi) construct targeting *mCherry* (control, UAS-*mCherry* RNAi) or *twinstar* (UAS-*tsr* RNAi), the only *DmCFL* gene. *MhcGal4>UAS-tsr* RNAi (*DmCFL* KD) larvae have decreased *DmCFL* RNA (27.9% of control level) and DmCFL protein (38.2% of control level) in larval body wall muscles (Fig. S1A-C). Levels of inactive phosphorylated DmCFL (p-DmCFL) were similarly reduced (32.1% of control level).

The *DmCFL* KD model showed a progressive muscle deterioration phenotype that we separate into three classes. Class 1 muscles retained sarcomeric actin organization in the muscle cell, class 2 muscles developed accumulation of actin at the cell poles with retained sarcomeric organization, and class 3 muscles lost overall muscle and sarcomeric integrity, in addition to having actin accumulations throughout the cell (Balakrishnan et al., 2020). Our data show a deterioration progression from class 1 into class 2 muscles, which, ultimately, give rise to class 3 muscles. At the final wandering third instar, all three muscle classes were present in an individual larva, with classes 2 and 3 predominating. In the present study, we analyzed ventral longitudinal muscles VL3 and VL4 (also known as muscles 6 and 7, respectively). These paired muscles are well-characterized, easily accessible, and share the same motor neuron innervation. We first performed a series of imaging and transcriptomic experiments to gain better understanding of the *DmCFL* KD phenotype.

First, to characterize the frequencies of the different muscle class combinations, we analyzed the F-actin sarcomeric structure at the wandering third instar stage based on phalloidin labeling. Despite sharing the same orientation, body position, and motor neuron, we found that these two muscles can be different classes, and thus, are not dependent on each other for their structural integrity (Fig. 1A). We considered the combination of VL3/4 classes as groups. Group 1 pairs had only class 1 muscles, while group 2 pairs had at least one class 2 muscle and group 3 pairs had at least one class 3 muscle. Most muscle pairs in wandering third instar larvae consist of two class 3 muscles (62%), yet even at this late stage some muscle pairs consist of two class 1 muscles (16%, Fig. 1B). We used confocal microscopy to image live larvae expressing a DmCFL::GFP protein, which showed that DmCFL localized to both the muscle sarcomere and myonuclei (Fig. 1C-E). Using immunofluorescence and structured illumination microscopy (SIM), we confirmed that DmCFL is present at the sarcomere Z-line and H-zone (Fig. 1F). Importantly, DmCFL is reduced at the sarcomere in *DmCFL* KD (Fig. 1F; Balakrishnan et al., 2020).

**Fig. 1.**
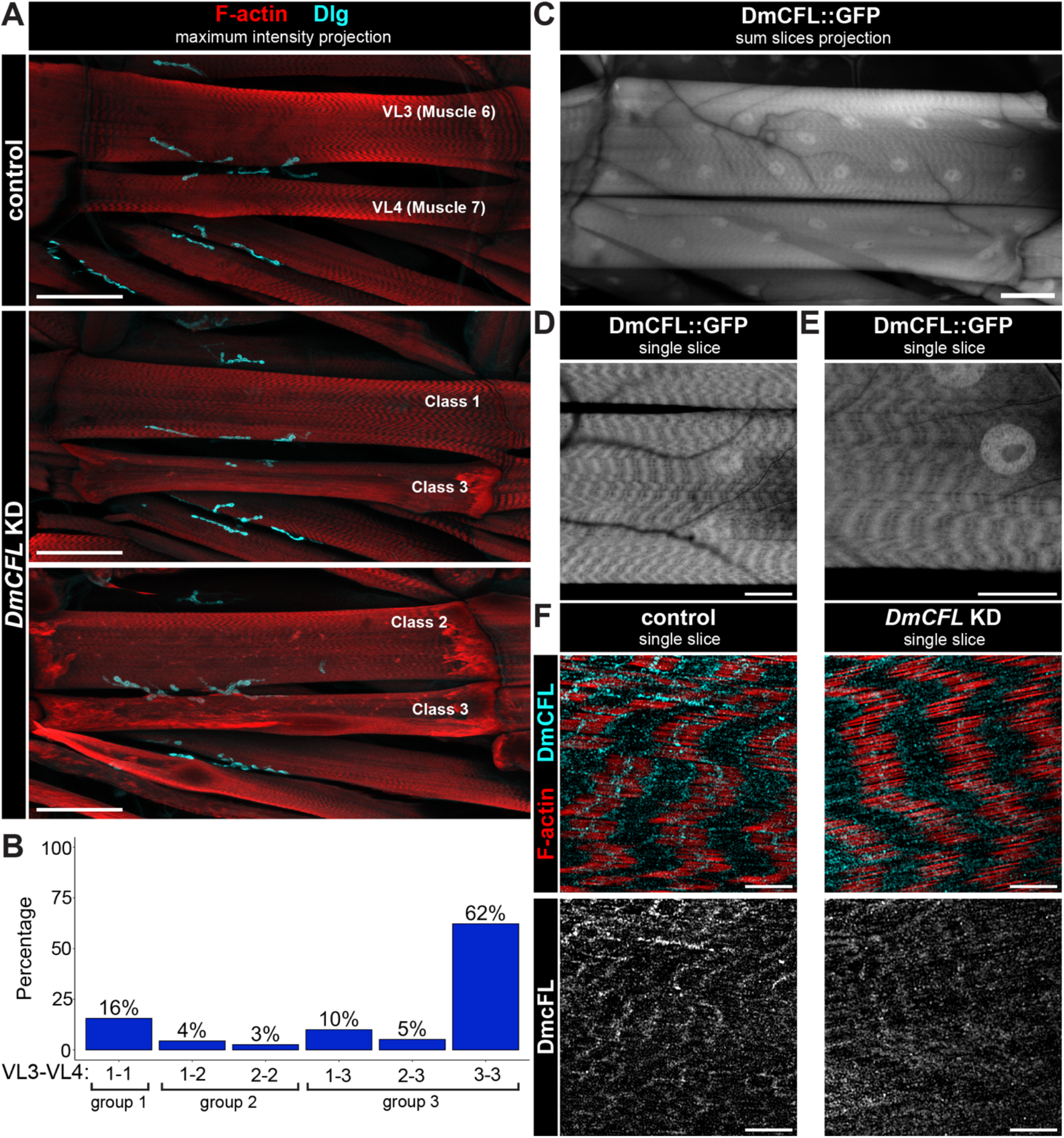
*DmCFL* KD reduction leads to muscle pairs of different deterioration classes. (A) Confocal images of larval ventral longitudinal (VL) muscles VL3 (6) and VL4 (7) labeled with phalloidin (red) and shared NMJ labeled with anti-Dlg (cyan) in control (top) and *DmCFL* KD (bottom) muscles. Muscles in the same pair (VL3 and VL4) can be of different phenotypic classes in *DmCFL* KD. Scale bar = 100 μm. (B) Quantification of the proportion of muscle class combinations (groups) in *DmCFL* KD larva (*n* = 45 larvae, 270 muscle pairs). (C) Confocal image of VL3-VL4 muscle pair expressing the protein trap DmCFL::GFP. Scale bar = 50 μm. (D,E) Magnified confocal images of DmCFL protein in the muscle sarcomeres of a live larva expressing the protein trap DmCFL::GFP. Scale bars = 25 μm. (F) Structured illumination microscopy (SIM) images of sarcomeres in control (left) and *DmCFL* KD (right) muscles labeled with phalloidin (red) and anti-DmCFL (cyan). Bottom panels show DmCFL expression alone (grayscale). Scale bar = 5 μm.

To quantifiably test the transcriptomic profile of *DmCFL* KD muscles, we dissected third instar larvae to produce muscle-enriched preparations for bulk RNA sequencing (Fig. 2; Fig. S2). Differential expression analysis using the DESeq2 package revealed a total of 1417 differentially-expressed genes with at least a two-fold change in *DmCFL* KD compared to control: 558 genes with decreased and 850 genes with increased expression (Fig. 2A; Fig. S2C-D). Reduced *DmCFL* expression was further confirmed in *DmCFL* KD muscle (Fig. S2B). RNA sequencing identified changes in genes related to actin binding or regulation. Overrepresentation analysis using the Gene Ontology (GO) biological processes gene sets showed that protein degradation processes were significantly reduced (Fig. 2B, Table S1; Ashburner et al., 2000; Gene Ontology Consortium et al., 2023). This finding was consistent with our previous work where increasing proteasomal activity in *DmCFL* KD larvae improved the muscle deterioration phenotype (Balakrishnan et al., 2020). Prominently, upregulated pathways were related to synaptic signaling (Fig. 2C, Table S1), due to increased expression of several genes acting at the neuromuscular junction (NMJ; Fig. 2D). These gene products have been previously reported to be functional on the presynaptic (i.e. motor neuron) and the postsynaptic (i.e. muscle) sides of the NMJ.

**Fig. 2.**
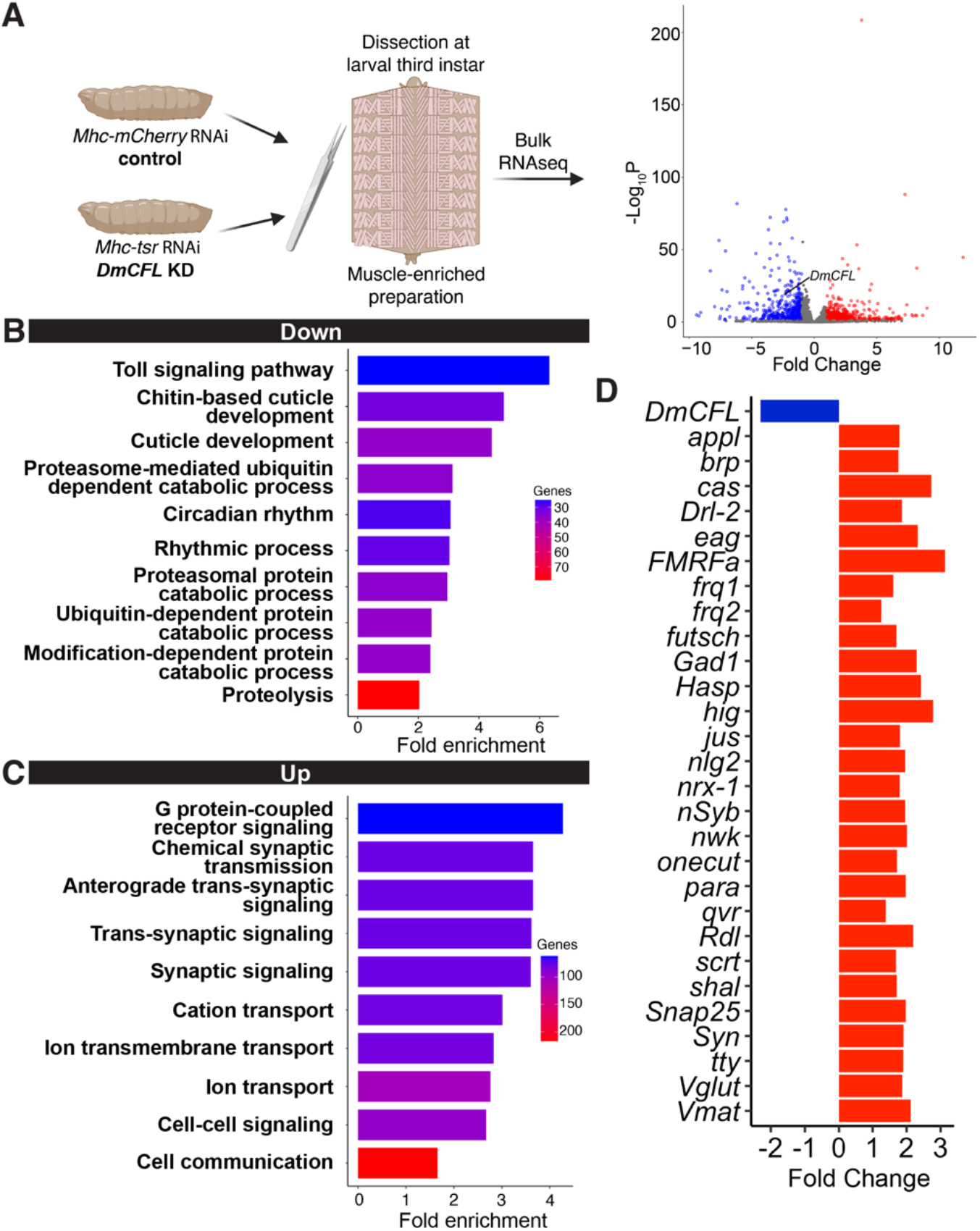
Transcriptomic analysis identifies changes at the neuromuscular junction (NMJ) in in *DmCFL* KD muscles. (A) Wandering third instar larvae with muscle *mCherry* RNAi (control) or *tsr* RNAi (*DmCFL* KD) were dissected, RNA extracted, and the RNA sequenced. Right: The volcano plot shows expression profiles in control and *DmCFL* KD with significantly upregulated (red) and downregulated (blue) genes marked. N = 3 replicates, each with *n =* 7-10 larvae per genotype. Diagram made using Biorender. (B) Gene ontology (GO) biological processes (BP) pathways identified as decreased significantly in *DmCFL* KD compared to control (false discovery rate 0.05, minimum fold change −2). (C) GO BP pathways identified as increased significantly in *DmCFL* KD compared to control (false discovery rate 0.05, minimum fold change 2). (D) Fold change of select genes related to the motor neuron and NMJ function.

Given these identified transcriptomic changes related to motor neuron-muscle communication at the NMJ, we next examined DmCFL and actin cytoskeleton in the postsynaptic domain.

### DmCFL protein is enriched at the muscle postsynapse and is reduced in *DmCFL* KD

The specialized postsynaptic muscle membrane, known as the subsynaptic reticulum (SSR), can be visualized using an antibody against the Discs-large (Dlg) scaffold protein (Lahey et al., 1994). The motor neuron presynaptic membrane can be visualized using anti-horseradish peroxidase (HRP; Jan and Jan, 1982). The SSR apposes the motor axonal endings (boutons), with distinctive Dlg rings around HRP signal (Fig. 3A). To test the presence of DmCFL at the postsynapse, both live and fixed imaging were used. For the following experiments, we focused our analyses on muscle pairs that had at least one class 1 or 2 muscle, since the postsynapse could not be reliably quantified in muscle pairs with both class 3 muscles due to their severe muscle deterioration.

**Fig. 3.**
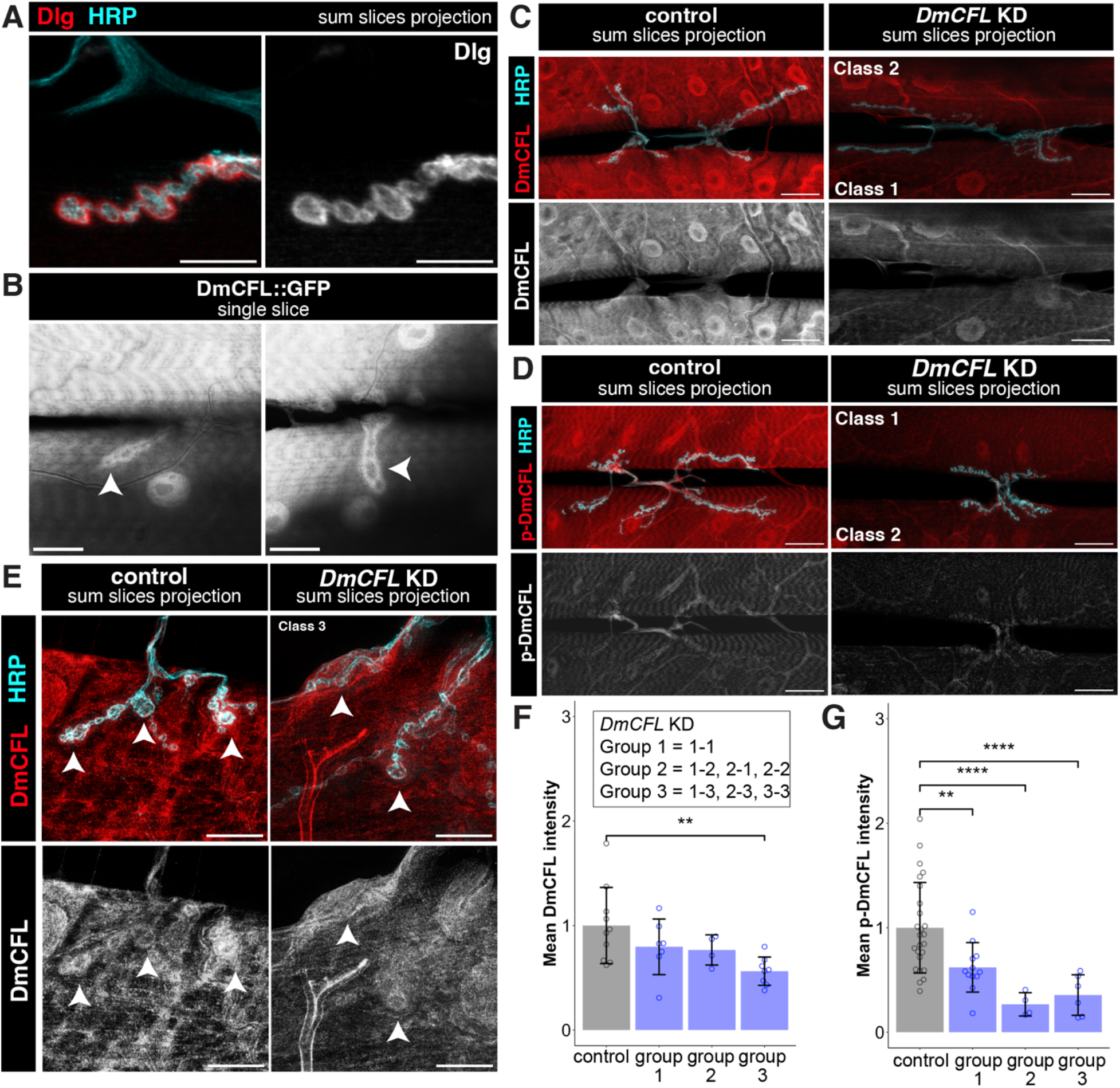
DmCFL localizes to the postsynapse and is reduced in KD muscles. (A) Left: Confocal images of NMJ labeled by postsynaptic membrane anti-Dlg (red) and presynaptic membrane labeled by anti-HRP (cyan). Specific *DmCFL* KD classes noted. Right: Dlg channel alone (grayscale). Scale bar = 10 μm. (B) Two representative confocal images of DmCFL protein at the postsynapse of the NMJ in live larvae expressing the protein trap DmCFL::GFP. Arrowheads indicate DmCFL at postsynaptic surrounding individual boutons. Scale bar = 20 μm. (C) Top: Confocal images of larval NMJ in control (left) and *DmCFL* KD (right) muscles labeled with anti-DmCFL (red) and anti-HRP (cyan). Bottom: DmCFL channel alone (grayscale). Scale bar = 25 μm. (D) Top: Confocal images of larval NMJ in control (left) and *DmCFL* KD (right) muscles labeled with p-DmCFL (red) and anti-HRP (cyan). Specific *DmCFL* KD classes noted. Bottom: p-DmCFL channel alone (grayscale). Scale bar = 25 μm. (E) Top: SIM images of NMJ boutons at the larval NMJ in control (left) and *DmCFL* KD (right) muscles labeled with anti-DmCFL (red) and anti-HRP (cyan). Bottom: DmCFL single channel (grayscale). Arrows indicate DmCFL at postsynaptic surrounding individual boutons. Scale bar = 10 μm. (F) Quantification of mean postsynaptic DmCFL intensity normalized to control (control 1 ± 0.365, *n =* 11 NMJs; overall *DmCFL* KD 0.692 ± 0.217, *n =* 19 NMJs). (G) Quantification of mean postsynaptic p-DmCFL intensity at postsynapse normalized to control (control 1 ± 0.433, *n =* 24 NMJs; overall *DmCFL* KD 0.484 ± 0.245, *n =* 22 NMJs). (F) Quantifications show mean ± SD with significance calculated by student’s t test (* p < 0.05, ** p < 0.01, **** p ≤ 0.0001).

Live imaging in DmCFL::GFP larvae showed the protein is enriched at the muscle SSR in a pattern consistent with the ring patterns of the postsynaptic density (Fig. 3B). In fixed samples, immunofluorescence and confocal microscopy showed that both total and inactive p-DmCFL are found at the postsynapse (Fig. 3C,D). However, we found that both DmCFL and p-DmCFL are reduced overall within *DmCFL* KD muscles. SIM imaging confirmed that DmCFL is found at the postsynapse in opposition to the boutons and that this local postsynaptic DmCFL is reduced in *DmCFL* KD muscles (Fig. 3E).

Given that the antibody against DmCFL also recognized DmCFL in the motor neuron, we sought to restrict our analysis to the postsynaptic domain while also minimizing signal from the underlying sarcomeres (approach described in detail in Fig. S3). We found that postsynaptic total DmCFL and p-DmCFL were both reduced in KD muscles: 69.2% and 48.4% of control levels, respectively (Fig. 3F,G).

Together, these data suggest that DmCFL and inactive p-DmCFL are enriched at the postsynaptic domain, and both are reduced in this region in in the *DmCFL* KD model.

### Postsynaptic actin organization is disrupted in *DmCFL* KD muscles

Regulation of F-actin by actin-binding proteins is important at the SSR membrane for proper cytoskeletal organization (Blunk et al., 2014; Pielage et al., 2006; Ramachandran et al., 2009; Wang et al., 2010). In the *DmCFL* KD model, we previously described aberrant F-actin accumulation at the fiber poles in class 2 and actin aggregates in class 3 muscles (Balakrishnan et al., 2020). Similarly, we now show that F-actin accumulates in the postsynaptic region in class 2 and 3 *DmCFL* KD muscles (Fig. 4A-B). In three-dimensions, the actin cytoskeleton formed a halo that enveloped the HRP-positive synaptic bouton, even if there were intact sarcomeres present deeper in a class 2 muscle (Movie 1). These data indicate that the F-actin network at the postsynaptic domain was affected prior to the complete muscle sarcomere deterioration seen in class 3 muscles. Postsynaptic F-actin accumulation occurred in both class 2 and 3 muscles independent of the class of its adjacent muscle in the VL3/4 muscle pair, suggesting that the effect is autonomous to muscles that have transitioned to a progressively deteriorated state. For example, a pair in which VL3 is class 1 and VL4 is class 3, only the class 3 muscle had aberrant postsynaptic F-actin accumulation. Therefore, the F-actin cytoskeleton at the muscle postsynapse is initially impacted and progressively deteriorates, similar to the deterioration seen in the sarcomeres in *DmCFL* KD muscles.

**Fig. 4.**
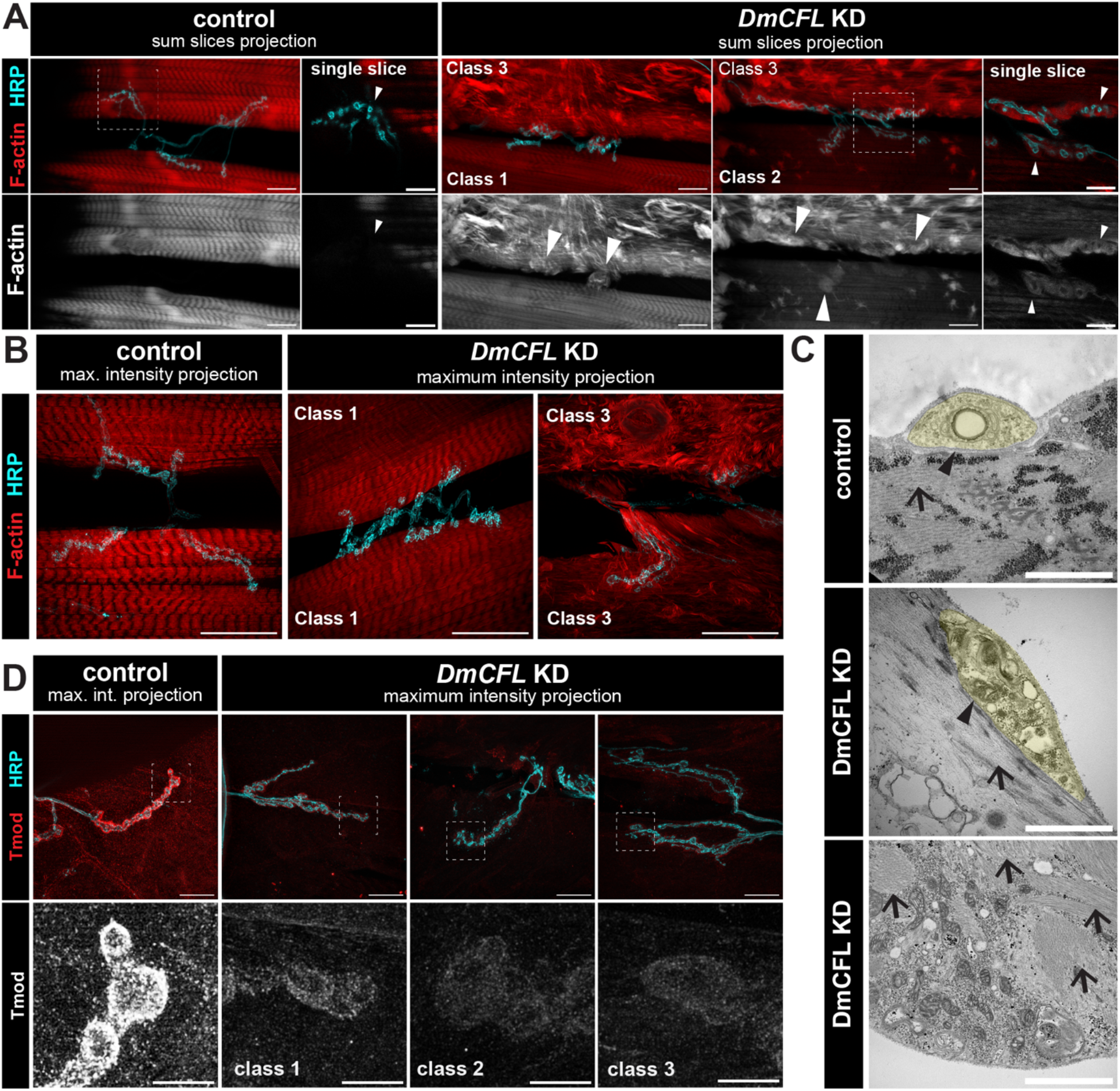
Actin and actin-binding proteins are disorganized at the postsynapse in *DmCFL* KD Class 2 and 3 muscles. (A) Top: Confocal images of larval NMJ in control (left) and *DmCFL* KD (right) muscles labeled with phalloidin (F-actin, red) and anti-HRP (cyan). Bottom: F-Actin single channel (grayscale). Arrowheads indicate accumulation of actin surrounding the boutons in *DmCFL* KD which is absent in control. Scale bar = 20 μm. Magnification of boxed area. Scale bar = 10 μm. (B) SIM images of larval NMJ in control (left) and *DmCFL* KD (right) muscles labeled with phalloidin (red) and anti-HRP (cyan). Specific muscle classes noted in DmCFL KD. Scale bar = 25 μm. (C) NMJ electron micrographs (sagittal sections) in control (top) and *DmCFL* KD (middle). Presynaptic boutons (pseudocolor in yellow) with muscle interface (arrowheads) and actin filaments (arrows). Bottom: disorganized actin filaments (arrows) throughout cytoplasm. Scale bar = 1 μm. (D) Top: SIM images of larval NMJ in control (left) and *DmCFL* KD (right) muscles of different classes labeled with anti-Tmod (red) and anti-HRP (cyan). Bottom: Tmod single channel (grayscale). Scale bar = 20 μm. Magnification shown below with scale bar = 5 μm.

Balakrishnan *et al*., showed that *DmCFL* KD class 3 muscles have protein aggregates containing actin and the Z-disc protein alpha-actinin, similar to nemaline bodies. By transmission electron microscopy (TEM), these structures were not as electron-dense as the rods seen in diseased human muscle (Balakrishnan et al., 2020). To test whether these nemaline body-like structures occur at the NMJ, we used TEM to examine the ultrastructure of the postsynaptic domain. Samples were dissected and sectioned longitudinally to reveal a sagittal view of the neuronal boutons and underlying muscle, including the postsynaptic actin network and sarcomeres. In control muscles, we observed actin filaments below the bouton organized parallel to each other (Fig. 4C). The postsynaptic actin organization below the bouton was lost in *DmCFL* KD. In addition, we found that there is a deformation of the synaptic cleft, the space between the bouton and the SSR membrane, in the *DmCFL* KD muscle. Importantly, the actin-capping protein tropomodulin (Tmod) was clearly present in the control, but it was greatly reduced in all *DmCFL* KD classes and became disorganized in classes 2 and 3 (Fig. 4D). Thus, the actin-binding protein Tmod did not display an accumulation pattern in *DmCFL* KD muscles, despite the observed actin accumulation in the postsynaptic domain.

As visualized by confocal microscopy and TEM, postsynaptic actin accumulates and is disorganized in *DmCFL* KD deteriorated muscles, surrounding the boutons in class 2 and 3 but not in class 1 muscles.

### Reduction of *DmCFL* in muscle affects postsynaptic morphology as muscles deteriorate

Genetic manipulations of other muscle actin-binding and regulatory proteins affect NMJ development leading to morphology abnormalities, such as reduced bouton number and Dlg disorganization (Lee and Schwarz, 2016; Pielage et al., 2006; Ramachandran et al., 2009; Xing et al., 2018). Thus, we sought to characterize NMJ morphology in *DmCFL* KD muscles.

The motor neuron properly innervated all *DmCFL* KD groups by the wandering third instar stage (Fig. 5A). The NMJ span along the muscle cell was unchanged (Fig. 5B). We did not observe any changes in the number of large Ib boutons in *DmCFL* KD muscles (Fig. 5C). To evaluate the postsynaptic SSR, we measured SSR area by taking the ratio of Dlg-positive area to the combined cell area of the muscle pair. Area quantification was done for group 1 or 2 VL3/4 muscle pairs, as cell area is difficult to quantify reliably when muscle cell integrity is severely compromised in group 3. We did not find a significant difference in SSR coverage when comparing the *DmCFL* KD to control (Fig. 5D). Morphologically, however, we detected differences in the postsynaptic organization: despite Dlg being properly organized around the boutons in class 1 KD muscles, it became increasingly disorganized in class 2 and 3 muscles, showing a more diffuse pattern (Fig. 5E). The disorganization in class 2 muscles was not enough to alter the mean SSR coverage compared to control. The range of disruption in Dlg organization over the three classes suggests that postsynaptic structural integrity deteriorates progressively, similar to defects seen in postsynaptic F-actin and the sarcomeres in the *DmCFL* KD model.

**Fig. 5.**
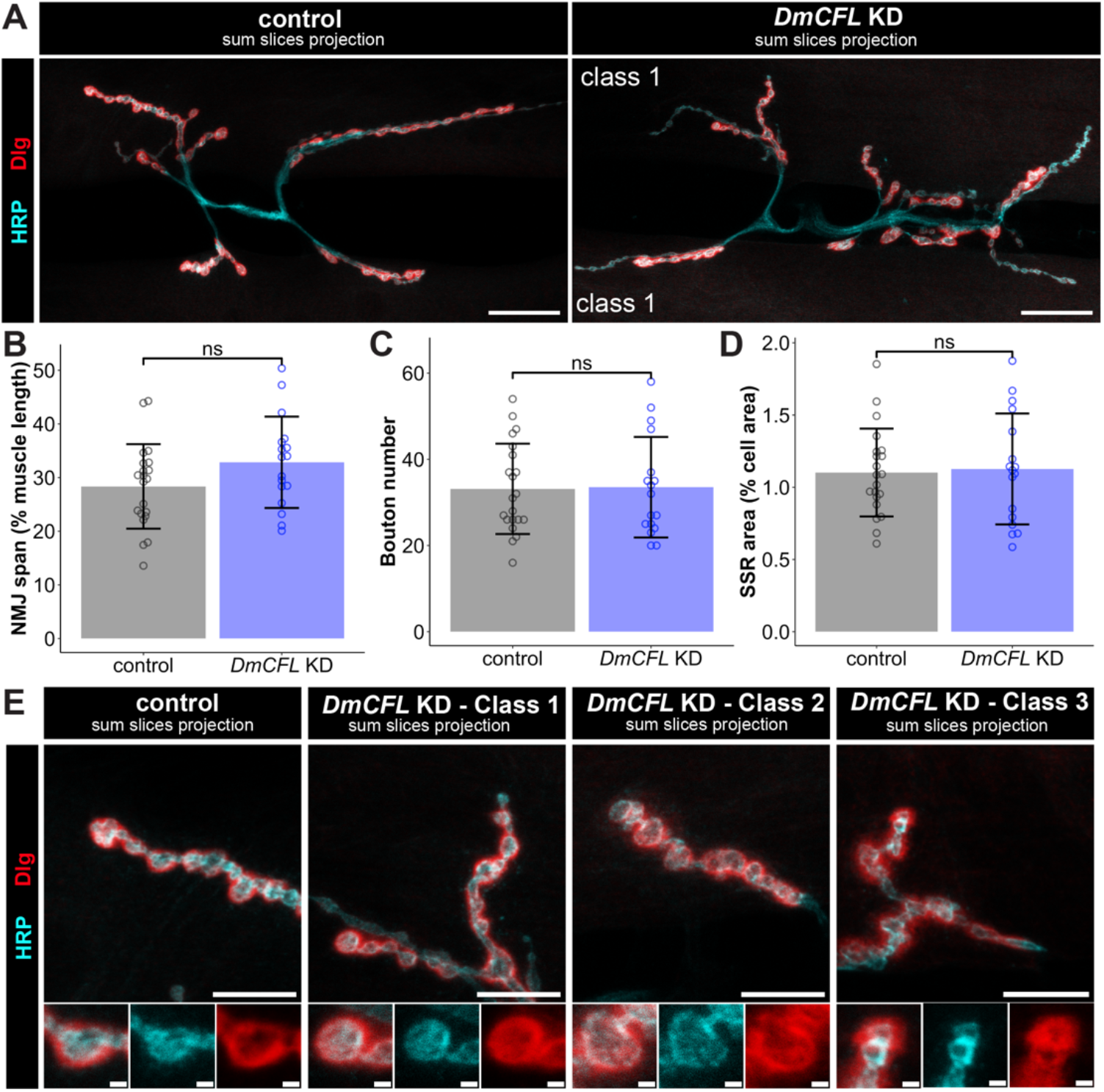
*DmCFL* KD affects postsynaptic morphology but not NMJ span or subsynaptic reticulum area. (A) Confocal images of larval NMJ in control (left) and *DmCFL* KD (right) muscles labeled with anti-Dlg (red) and anti-HRP (cyan). Scale bar = 25 μm. (B) Quantification of NMJ span, defined as NMJ length normalized to cell length reported as percent of muscle length (control 28.35 ± 7.87%, *n =* 21 NMJs; *DmCFL* KD 32.84 ± 8.51%, *n =* 17 NMJs). (C) Quantification of total Ib (large) bouton number (control 33 ± 11, *n =* 21 NMJs; *DmCFL* KD 34 ± 12, *n =* 17 NMJs). (D) Quantification of subsynaptic reticulum (SSR) area, defined as Dlg-positive area normalized to cell area (control 1.10 ± 0.30, *n =* 21 NMJs; *DmCFL* KD 1.13 ± 0.38, *n =* 17 NMJs). (E) Top: Confocal images of larval NMJ in control (left) and *DmCFL* KD (right) class 1, class 2 and class 3 muscles labeled with anti-Dlg (red) and anti-HRP (cyan). Scale bar = 10 μm Bottom: Panels show individual boutons (merge and individual channels). Scale bar = 1 μm. Quantifications show mean ± SD with significance calculated by student’s *t* test (ns = not significant, *p* > 0.05).

Together, these data indicate normal presynaptic innervation when *DmCFL* is reduced in the muscle, whereas the postsynaptic architecture became progressively disorganized as the muscle deteriorates over time in the *DmCFL* KD model.

### Presence of presynaptic Brp and postsynaptic GluRIIC is unaffected by muscle *DmCFL* KD

Excitation-contraction coupling is the process by which synaptic signaling from the motor neuron bouton is communicated to the muscle to drive contraction. In *Drosophila*, this neurotransmission relies on glutamatergic signaling, while in vertebrates excitation occurs via acetylcholine. At the presynaptic active zone, a consolidation of proteins, including the core active zone scaffold Bruchpilot (Brp), orchestrate synaptic vesicle docking and release of glutamate (Wagh et al., 2006). In the postsynaptic SSR membrane, the excitatory neurotransmitter binds to glutamate receptors (GluR). These receptors are heterotetramers, with essential subunits GluRIIC, D, and E, and an optional fourth subunit of either GluRIIA or GluRIIB (DiAntonio, 2006). To test whether there were changes in NMJ signaling machinery are affected in *DmCFL* KD muscles, we analyzed Brp and the essential GluRIIC subunit.

While our RNA sequencing results indicated an over three-fold increase in *brp* gene expression (Fig. 2D), we found that Brp protein in NMJ presynaptic boutons was not detectably changed in *DmCFL* KD NMJs compared to control (Fig. 6A-B). Likewise, the postsynaptic GluRIIC protein level was unchanged in *DmCFL* KD muscles, suggesting that there was no change in the total amount of glutamate receptors in the postsynaptic membrane (Fig. 6C). No gross differences were detected in the apposition of GluR to Brp (Fig. 6A).

**Fig. 6.**
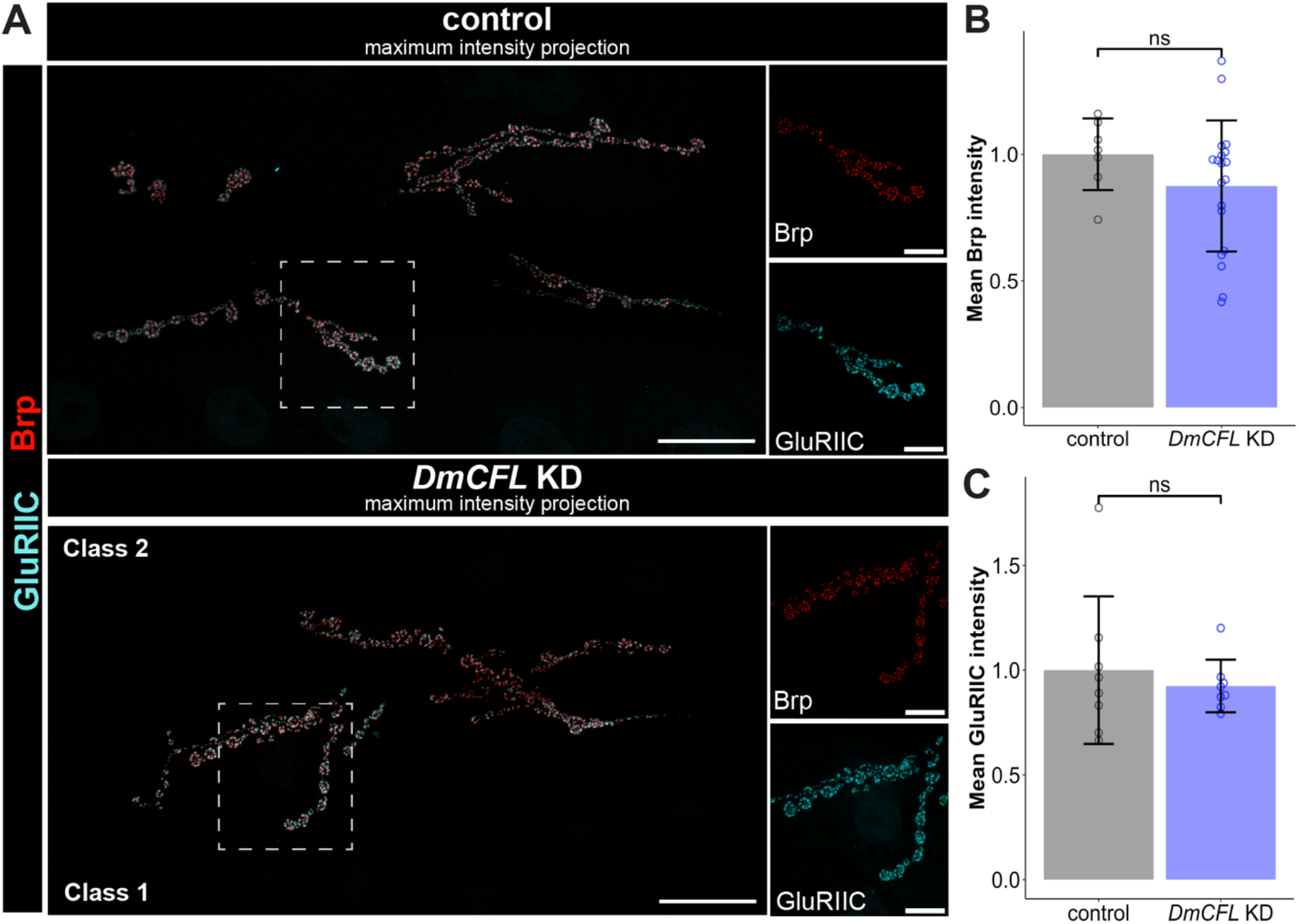
Presynaptic Brp and postsynaptic GluRIIC unchanged with *DmCFL* KD. (A) Left: Confocal images of control (top) and DmCFL KD (bottom) labeled with presynaptic anti-Brp (red) and postsynaptic anti-GluRIIC (cyan). Right: magnification of boxed areas, with each stain shown separately. Scale bar = 25 μm, magnification scale bar = 10 μm. (B) Quantification of mean Brp intensity normalized to control (control 1 ± 0.14, *n =* 8 NMJs; *DmCFL* KD 0.87 ± 0.26, *n =* 19 NMJs of all groups). (C) Quantification of mean GluRIIC intensity normalized to control (control 1 ± 0.35, *n =* 9 NMJs; *DmCFL* KD 0.92 ± 0.13, *n =* 8 NMJs of all groups). Quantifications show mean ± SD with significance calculated by student’s *t* test (ns = not significant, *p* > 0.05).

These results suggest that the *DmCFL* KD does not affect protein levels nor organization of presynaptic active zone Brp or the postsynaptic GluRs at the NMJ, despite highly increased RNA expression of NMJ genes being identified by transcriptomic analysis. Given that there is diversity in GluR subunit composition that impacts GluR channel properties, we pursued further synaptic analyses in *DmCFL* KD.

### *DmCFL* KD disrupts glutamate receptor composition and neurotransmission

A previous screen examining lethal mutants qualitatively identified various cytoskeleton-related genes that specifically affect GluRIIA subunit levels detected by immunofluorescence (Liebl and Featherstone, 2005). GluRIIA-containing receptors in the postsynaptic domain are selectively regulated by Coracle via the cortical actin cytoskeleton (Chen et al., 2005; Song et al., 2022). Moreover, actin monomer-binding protein twinfilin mutants have decreased abundance of GluRIIA-containing receptors at the NMJ (Wang et al., 2010). Since DmCFL is an actin-binding and severing protein, we hypothesized that muscle *DmCFL* KD could selectively affect GluRIIA levels.

To test this hypothesis, we quantified GluRIIA using immunofluorescence, and we found that there was less GluRIIA in *DmCFL* KD muscles compared to control (66%; Fig. 7A). Stratifying the data by muscle pair class combination showed that GluRIIA is only decreased in group 2 or 3 muscle (Fig. 7B). SIM imaging revealed that glutamate receptors containing GluRIIA were properly located at the NMJ postsynaptic domain in class 1 muscles, but the field of these receptors was expanded and mislocalized at some postsynaptic densities in class 2 and class 3 muscles (Fig. 7C). GluRIIA, unlike other GluR subunits, is anchored to the actin cytoskeleton by Coracle (Chen et al., 2005). Consistent with the Coracle regulatory mechanism, we found that *DmCFL* KD muscles have reduced Coracle in the postsynaptic domain, suggesting a possible mechanism for altered GluRIIA localization at the NMJs of *DmCFL* KD muscles (Fig. 7D).

**Fig. 7.**
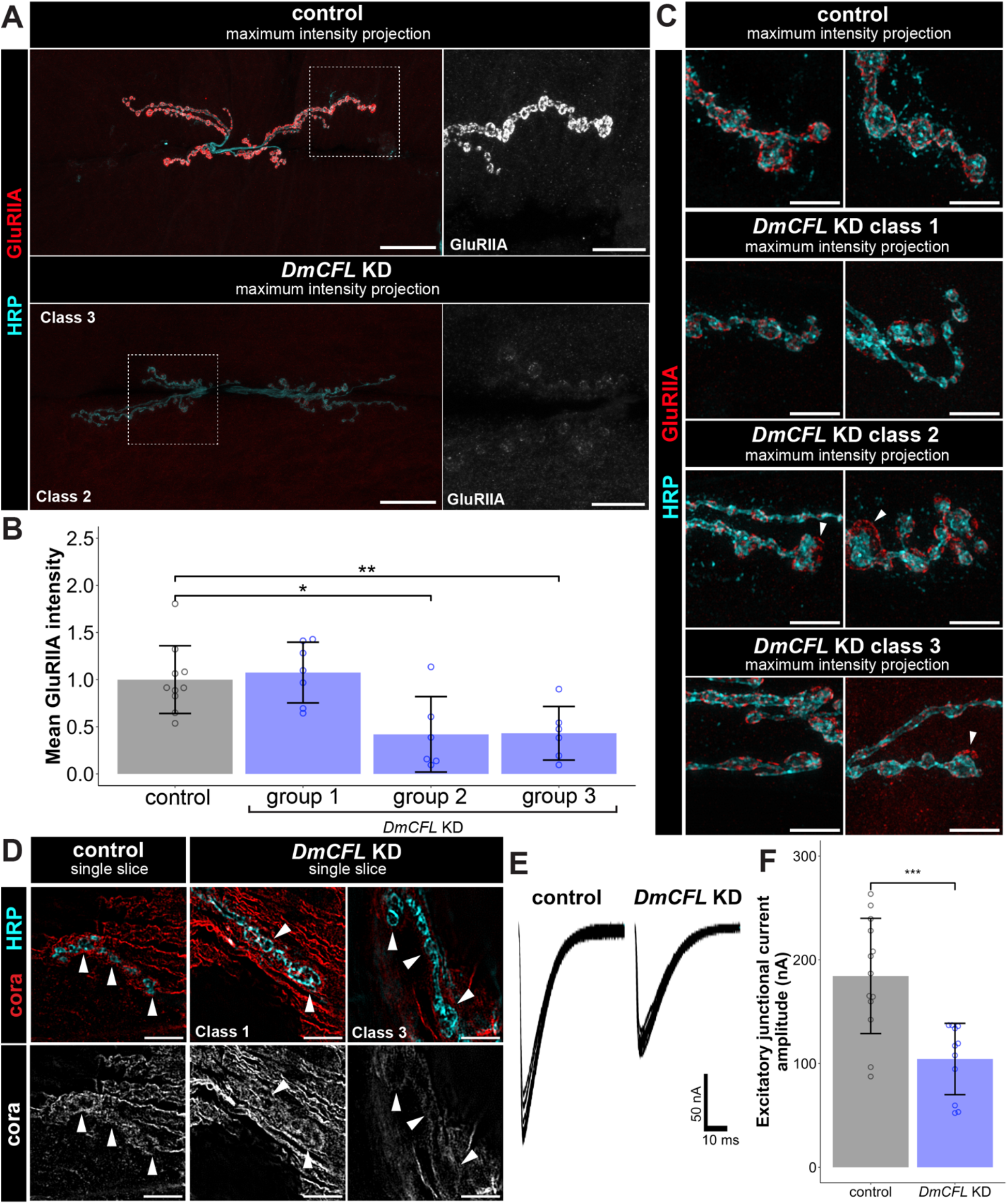
*DmCFL* KD reduced NMJ GluRIIA levels and neurotransmission strength. (A) Left: Confocal images labeled with anti-GluRIIA (red) and anti-HRP (cyan) Top, Control; Bottom, *DmCFL* KD. Scale bar = 25 μm. Right: magnification of boxed areas, showing GluRIIA channel alone (grayscale). Scale bar = 10 μm. (B) Quantification of mean GluRIIA intensity normalized to control by muscle group (control 1 ± 0.36, n = 10 NMJs; *DmCFL* KD 0.66 ± 0.45, n = 19 NMJs). (C) SIM images of boutons labeled with anti-GluRIIA (red) and HRP (cyan). Top: Control; Bottom: *DmCFL* KD muscle classes. Arrowheads indicate GluRIIA fields that do not overlap with HRP. Scale bar = 5 μm. (D) Top: SIM images labeled with anti-Coracle (red) and anti-HRP (cyan) in control (left) and DmCFL KD (right) muscle classes. Bottom: Coracle single channel (grayscale). Scale bar = 5 μm. (E) Example evoked excitatory junctional current (EJC) traces showing ten consecutive evoked traces at 0.2 Hz. (F) Quantification of EJC amplitude from individual NMJs (control 184.4 ± 55.52 nA, n = 11 NMJs; *DmCFL* KD 104.4 ± 034.37 nA, n = 13 NMJs). Quantification shows mean ± SD with significance calculated by student’s t test (* p ≤ 0.05, ** p ≤ 0.01; *** p ≤ 0.001).

GluRIIA-containing receptors mediate the strength of the muscle postsynaptic response, with reduced GluRIIA selectively impairing NMJ neurotransmission strength (DiAntonio et al., 1999; Petersen et al., 1997). To examine neurotransmission in *DmCFL* KD muscle, we used two-electrode voltage-clamp (TEVC) recordings to measure glutamate release and GluR activation (Leahy et al., 2023). The motor neuron was stimulated at suprathreshold voltages with a glass suction electrode at 0.2 Hz, and 10 consecutive evoked excitatory junction (EJC) traces were recorded (Fig. 7E) and then averaged to calculate the mean neurotransmission strength in the *DmCFL* KD compared to control (Fig. 7F). To get a varied sample of the electrophysiological profile of *DmCFL* KD muscles, a mix of the different muscle class combinations were tested by TEVC in a blinded configuration. Compared to control, the overall EJC amplitude of *DmCFL* KD muscles was significantly lower (about 60% reduced; Fig. 7E,F). Interestingly, all *DmCFL* KD muscles had a decreased EJC amplitude and, therefore, similarly reduced neurotransmission strength. This indicates that NMJ function was impaired independent of muscle degeneration when DmCFL levels are reduced.

These findings indicate that *DmCFL* KD NMJs have decreased levels of GluRIIA-containing receptors, with mislocalization of the receptors into extra-synaptic muscle regions and a concomitant reduction in synaptic transmission amplitude shown by direct voltage-clamp electrophysiology recording.

## Discussion

Cofilin-2, an actin-severing protein, has been linked to NM. While NM is often characterized as a disease of the sarcomere, other actin dependent structures and functions in the muscle have not been investigated. Consistent with its known role in human NM, we have shown that DmCFL plays a critical role in sarcomerogenesis and muscle maintenance (Balakrishnan et al., 2020). Here, we investigated an additional role of CFL in regulating actin at the NMJ. We show that DmCFL is located at the postsynaptic domain and that loss of DmCFL results in buildup of postsynaptic F-actin, disorganization of SSR membrane proteins, and mislocalization of the key GluRIIA glutamate receptor subunit. Consistent with these findings, we see reduced NMJ neurotransmission by electrophysiology. Interestingly, these defects occur prior to the onset of advanced sarcomeric decline. We propose that DmCFL is required for proper postsynaptic actin dynamics, and, in its absence, the SSR membrane and its components become progressively disorganized.

As a first step to study the role of DmCFL at the *Drosophila* NMJ and to determine its postsynaptic localization and reduction in *DmCFL* KD muscles, we visualized DmCFL in live and fixed tissues using different approaches. DmCFL is found at the postsynapse, which can only be appreciated by examining thin sections and suggests that its localization is fairly restricted in space. We used an innovative three-dimensional approach to quantify proteins at the postsynaptic domain to distinguish from the high levels of DmCFL in the underlying sarcomere. This approach could be applied to other studies of the postsynapse. Postsynaptic quantification, in addition to Western blot of overall muscle levels, revealed that total DmCFL and inactive p-DmCFL are reduced in *DmCFL* KD. Similarly, phosphorylated cofilin-2 is absent in the muscles of an NM patient harboring a *CFL2* mutation (Agrawal et al., 2007).

In humans, mice, and *Drosophila* with muscle cofilin defects, muscle structure progressively deteriorates and function declines. Progressive muscle weakness is reported in *CFL2* NM cases with some patients showing a deterioration after early childhood and others having severe symptoms present at birth (Agrawal et al., 2007; Fattori et al., 2018; Ockeloen et al., 2012; Ong et al., 2014). Cfl2 mouse models also show a progressive decrease in muscle function and an increase in nemaline rods seen on histology (Agrawal et al., 2012; Gurniak et al., 2014; Mohri et al., 2019; Rosen et al., 2020). Even in chimeric mice where only some muscle fibers harbor *Cfl2* mutations, the mutant fibers deteriorate, indicating that health of surrounding cells does not impact muscle disease progression (Mohri et al., 2019). In *Drosophila*, *DmCFL* is knocked down in all larval muscles, yet the progression of the deterioration phenotype does not occur in a coordinated fashion across all muscles. We show that the pair of ventral longitudinal muscles VL3 and 4, which are innervated by the same branch of intersegmental nerve B, do not necessarily exhibit the same extent of deterioration by the end of larval development, as evidenced by the existence of various groups of classes. This finding mirrors the progressive deterioration seen in the chimeric *Cfl2* mice, since cofilin-2 is only affected in select muscles rather than in all tissues like in other *Cfl2* mouse models.

Our work demonstrates that both postsynaptic actin and SSR membrane organization deteriorate progressively in *DmCFL* KD muscles. In vertebrates, actin isoforms have been defined based on their expression: skeletal muscle (α_sk_ and α_ca_), smooth muscle (α_sm_ and γ_sm_) and cytoplasmic (β_cyto_ and γ_cyto_) [reviewed by Perrin and Ervasti, 2010; Lubit and Schwartz, 1980]. Despite the sarcomeric ⍺-actin isoforms being predominant in skeletal muscle, the non-sarcomeric γ-actin networks play myriad roles in the muscle cell. This isoform is needed for anchoring myofibrils to the muscle membrane, localization of organelles such as the mitochondria and sarcoplasmic reticulum, and as part of the Z-disc (Craig and Pardo, 1983; Gokhin and Fowler, 2011; Nakata et al., 2001; Papponen et al., 2009; Pardo et al., 1983; Rybakova et al., 2000). Additionally, γ_cyto_ is only needed for proper maintenance of the cytoskeleton in muscle but not its development; nevertheless, its absence does lead to a phenotype similar to centronuclear myopathy (Belyantseva et al., 2009; Sonnemann et al., 2006). Given that there are different pools of actin in the muscle cell, cofilin may play a role at the sarcomere engaging with ⍺-actin and at other locales—like the NMJ—by regulating γ_cyto_ actin. The fact that actin is critical in the muscle for more than sarcomere structure invites further inquiry into how affecting non-sarcomeric actin contributes to NM and defects at the postsynapse.

In addition, there is a greater increase in F-actin than G-actin seen in the muscles of a *Cfl2* knockout mouse model (Agrawal et al., 2012). The postsynaptic actin accumulations in the *DmCFL* KD appear as filamentous swirls surrounding the motor neuron boutons, similar to the structures reported in the *act^E84K^* mutant that affects the actin isoform Act57B and the *twf^110^* mutant that affects actin-severing protein twinfilin (Blunk et al., 2014; Wang et al., 2010). We found that defects in F-actin at the NMJ in class 2 muscles occurred before sarcomeric deterioration deeper in the muscle, indicating that it is unlikely that sarcomeric actin is the only source of actin accumulating at the NMJ. F-actin accumulations surround only some boutons in class 2 muscles, while all boutons are surrounded by actin in class 3 muscles, confirming that the deterioration is progressive. It is possible that the postsynaptic deterioration may also be progressive in human NM patients, and that accumulations may not be as obvious as the rods seen emanating from the sarcomere because they do not include Z-disc proteins often examined in patient samples. Even using TEM, we found disorganized actin filaments in the postsynaptic region of *DmCFL* KD muscle which is reminiscent of the cytoplasmic actin disorganization shown in the biopsy from one *CFL2* NM patient (Fattori et al., 2018). We find that actin-binding protein Tmod is reduced at the *DmCFL* KD postsynapse as well, which may be due to sequestering of Tmod at the cell poles as we have previously reported (Balakrishnan et al., 2020).

Several *Drosophila* models have shown that the actin cytoskeleton is important for proper SSR formation (Pielage et al., 2006; Ramachandran et al., 2009; Wang et al., 2011). While the postsynaptic membrane forms properly initially, the expansion of Dlg at the postsynapse in *DmCFL* KD class 3 muscles suggests that actin dynamics—regulated by DmCFL severing—are important in maintaining the SSR. We did not see an increase in postsynaptic SSR area in group 1 and group 2 pairs likely because only some boutons in class 2 muscles have affected Dlg organization. Dlg is thought to not interact directly with actin, instead being part of a complex with other actin-binding proteins, including adducin/Hts and spectrin (Wang et al., 2014). Thus, Dlg disorganization at the *DmCFL* KD postsynapse in the most deteriorated muscles is likely due to an effect on the overall complex, rather than the actin accumulation affecting Dlg directly. *Cfl2* mouse studies report no obvious NMJ defects; it is possible that a subtle change on the postsynaptic membrane would not be appreciated when only visualizing the motor neuron and acetylcholine receptors (Agrawal et al., 2012; Gurniak et al., 2014). Nevertheless, some reports from human NM patient biopsies imaged by TEM report alterations at the postsynapse, including collapsed or dilated primary and secondary synaptic clefts (Fukuhara et al., 1978; Heffernan et al., 1968; Karpati et al., 1971). We see a collapse of the synaptic cleft via TEM in the *DmCFL* KD model. Together, these data suggest that maintaining a certain amount and/or dynamic state of postsynaptic actin is important for retaining SSR structure, and, thus, the increase in actin due to cofilin reduction in the fly model leads to progressive structural deterioration.

Cofilin and the actin cytoskeleton have been implicated in the trafficking of ionotropic AMPA receptors (AMPAR) to dendritic membranes in vertebrates. In fact, it has been suggested that ADF/cofilin activity is needed both for clearing actin that may be in the way of AMPAR insertion and for regulating the new filaments that guide the receptors to the membrane (Gu et al., 2010; Zhou et al., 2001). Experimentally, an increase of ADF/cofilin in hippocampal neuron culture led to increased addition of AMPAR to dendritic spines while a decrease resulted in their removal (Gu et al., 2010). Work in a *Xenopus* culture model found that actin is more dynamic near the acetylcholine receptors (AchR) at the membrane compared to the actin within the myofibrils, and that a balance of active and inactive cofilin near the membrane is required for their proper addition (Lee et al., 2009). Data from *Drosophila* shows that pharmacological stabilization of actin at the NMJ reduces the GluRIIA cluster size (Chen et al., 2005). From the current studies in mouse, it is unclear if there is a reduction in AChR levels in *Cfl2* knockout mice. Our *DmCFL* KD experiments indicate that there is not a reduction in the levels of total glutamate receptors as indicated by the GluRIIC subunit present in all receptors, but there is reduced GluRIIA. Using SIM, we show that in class 2 and 3 muscles GluRIIA-containing receptors are being pulled away from or are not properly delivered to the postsynaptic density directly opposing the motor neuron. Coracle links the GluRIIA subunit to the actin cytoskeleton, and its loss reduces levels of GluRIIA (Chen et al., 2005). We show that there are reduced Coracle levels at the *DmCFL* KD postsynapse, which is consistent with the reduction in GluRIIA. A similar decrease in GluRIIA and Coracle levels is seen in a mutant of the actin-binding protein twinfilin (Wang et al., 2010). Live imaging experiments have revealed that glutamate receptors enter the postsynaptic density from pools around the plasma membrane; it is possible that without a dynamic actin cytoskeleton, GluRIIA receptors are not transported to the postsynapse (Rasse et al., 2005).

Our electrophysiological profile of *DmCFL* KD muscles is consistent with decreased levels of GluRIIA at the postsynapse. Electromyogram (EMG) studies of NM patients often result in a myopathic pattern with low amplitudes. However, one longitudinal study of 13 NM patients found that after age nine EMG studies of distal muscles began to show neuropathic changes in addition to myopathic changes (Wallgren-Pettersson et al., 1989). This suggests that as the disease progresses there is degenerative changes of the motor units with some patients reported to have lacked motor units; it is also possible that reinnervation by adjacent motor units occurs to result in higher amplitude potentials. One case report speculated that there was a disruption in innervation which could contribute to the formation of rods in the extrafusal muscle fibers (Karpati et al., 1971). Of note, we did not appreciate any defect in axon pathfinding to its VL3/4 muscle target nor axon structure at the level of the presynaptic active zones; however, the upregulation in presynaptic genes identified by RNA sequencing may suggest that an attempt by the motor neuron to compensate does occur at the final stage of deterioration in the *DmCFL* KD model. We are unable to further investigate this possibility since the larvae begin the process of pupation after the third-instar wandering stage.

While our study focuses on a cofilin NM model, the patient case reports suggest that NMJ defects may be found in NM more broadly. Future studies should examine NM patient biopsy and electrophysiological results with more consideration for the specific gene mutated, which was not possible in previous decades. The *DmCFL* KD model should encourage further study into the role of the NMJ in NM disease progression and potential therapeutics (Fisher et al., 2022). There are reports from individual NM patients that use of acetylcholinesterase inhibitors can lead to clinical improvement (Natera-de Benito et al., 2016). One possibility for the improvement is that increased presence of acetylcholine in the synaptic cleft increases the probability that the neurotransmitter will find receptor along the simplified postsynaptic membrane, thereby increasing NMJ signal transmission. New acetylcholinesterase inhibitors, such as C-547, that are more specific for the NMJ region show promise as they would limit off-target side effects of treatment (Petrov et al., 2018). Further study is needed into which pharmacologic treatments may work for NM. The fact that the *Drosophila* NMJ is a glutamatergic synapse poses a challenge in testing potential treatments that impact acetylcholine; however, fly models allow a simple system to further interrogate the basic science underpinnings of disease mechanism. Our findings suggest that NMJ defects are seen when muscle cofilin is reduced, which brings forth the alternative that druggable targets of actin-binding proteins could be screened in the fly. In addition, we observed NMJ structural deterioration in the most affected muscles, and thus EMG in tandem with nerve conduction studies and biopsies from older *CFL2* NM patients would be informative about disease progression and health of the NMJ.

In conclusion, our data point to changes at the NMJ, specifically the postsynapse, as being a part of the deterioration that results from affecting muscle cofilin levels. Affecting non-sarcomeric actin dynamics may affect other muscle functions that then contribute to the progression of muscle deterioration seen in NM, opening new avenues for understanding and ameliorating the condition.

## Materials and Methods

### *Drosophila* husbandry, stocks, and crosses

All stocks and crosses were raised on standard cornmeal at 25°C on a 12-hour light/12-hour dark light cycle under humidity control. All experiments were done in wandering L3 larvae of both sexes reared at 25°C. *DmCFL* was knocked down specifically in muscle, as done in our previous study (Balakrishnan et al., 2020), by leveraging the Gal4-UAS system (Brand and Perrimon, 1993). The muscle-specific driver *Mhc*-Gal4 (BDSC #67044) was used to drive UAS-*mCherry* RNAi (for control, BDSC #35785) or UAS-*tsr* TRiP RNAi (for *DmCFL* knockdown, BDSC #65055) generated by the Transgenic RNAi Project (TRiP; Perkins et al., 2015). Live imaging experiments were done with larvae expressing tsr::GFP (ZCL2393, Kyoto Stock Center DGRC #110875) generated by the FlyTrap: GFP Protein Trap Database (Morin et al., 2001).

### RNA-sequencing analysis

RNA-sequencing of control and *DmCFL* KD larvae as described previously (Zapater I Morales et al., 2023). Eight to ten late third instar larvae of each genotype were dissected in triplicate. Read counts data were assessed and plotted with the integrated Differential Expression and Pathway analysis (iDEP) web application (versions 0.96 and 1.1; Ge et al., 2018). DESeq2 analysis was performed with the following parameters: false discovery rate of 0.05 and minimum two-fold change (Love et al., 2014). Overrepresentation analysis was done using the Gene Set Enrichment Analysis (GSEA) method to identify the top 20 enriched pathways defined by Gene Ontology (GO) Biological Processes (BP) gene sets when considering the top 2000 genes at a false discovery rate of 0.05.

### Western blot

Five to ten wandering late third-instar larvae were dissected in HL3.1 buffer as has been previously described in (Brent et al., 2009) to produce muscle-enriched preparations, which were then lysed in larval lysis buffer (50 mM HEPES [pH 7.5], 150 mM NaCl, 0.5% NP40, 0.1% SDS) supplemented with cOmplete mini protease inhibitor cocktail (Roche, #11836153001) and PhosSTOP phosphatase inhibitor cocktail (Roche, #4906837001). Ten micrograms from control and *DmCFL* KD lysates were run on a 12.5% polyacrylamide gel, then transferred to a nitrocellulose membrane (Thermo Fisher Scientific, PI88018). Blocking was done in 5% milk or 5% bovine serum albumin (BSA) in TBST (Tris-buffered saline+0.1% Tween) for 1 hour at room temperature. Primary antibody staining was performed in Stamina Antibody Dilution Buffer (KindleBiosciences, R2004) overnight at room temperature using 1:1000 for rabbit anti-Twinstar (DmCFL, gift from Tadashi Uemura); 1:1000 for rabbit anti-phospho-Twinstar (p-DmCFL, gift from Tadashi Uemura); and 1:1000 for mouse anti-GAPDH (Abcam, #ab9484). Secondary antibody incubation was performed in 5% milk for 1 hour at room temperature using 1:5000 for anti-rabbit-HRP (Jackson Immunoresearch, 711-035-152) or anti-mouse-HRP (Jackson Immunoresearch, 715-035-151). Blot was imaged using the KwikQuant Imager (Kindle Biosciences, D1001) using 1-Shot Digital-ECL (Kindle Biosciences, R1003) and intensities were quantified using FIJI Gel Analyzer tool (NIH). Protein expression was normalized to GAPDH loading control within each sample and repeated in triplicate.

### Quantitative polymerase chain reaction (q-PCR)

Total RNA was extracted from ten late third-instar wandering larvae muscle-enriched preparations (dissected as described in western blot methods) using TRIzol reagent (ThermoFisher #15596026) and was subsequently cleaned using the TURBO DNA-free Kit (Ambion, AM1907). Reverse transcription was done to synthesize cDNA using the SuperScript III First-Strand Synthesis System for RT-PCR (Invitrogen #18080-051) kit. PCR reactions were run on the CFX Opus 96 Real-Time PCR System using SYBR Select Master Mix for CFX (Applied Biosystems #4472937) in biological and technical triplicate. Primers used for tsr: (forward) GCTCTCAAGAAGTCGCTCGT; (reverse) GCAATGCACAGTGCTCGTAC. The delta-delta Ct method was used to calculate fold changes with Rpl32 as the normalization control. Reported values represent the log_2_ fold change of the gene in *DmCFL* KD compared to expression in control samples.

### Dissection and immunostaining

Third-instar larvae at the wandering stage were dissected as previously described in (Azevedo et al., 2016; Brent et al., 2009) to expose the body wall muscles. Fixation was done using 4% paraformaldehyde in HL3.1 buffer for 15 minutes at room temperature for all antibodies except GluRIIA for which Bouin’s fixative was used for 5 min at room temperature. Samples were blocked in BSA-PBT (PBS supplemented with 0.1% BSA and 0.3% Triton X-100) for 30 minutes at room temperature, then incubated with primary antibody overnight at 4°C and subsequently washed in BSA-PBT. Samples were then incubated with Alexa Fluor-conjugated secondary antibodies, phalloidin and goat Alexa-647 conjugated horseradish peroxidase (HRP; Jackson Immunoresearch cat no. 123-605-021) at a concentration of 1:400. Alexa Fluor 555 conjugates secondary antibodies were used for all intensity quantifications. Final washes were done in PBT prior to mounting in Prolong Gold (Invitrogen, P36930). All slides were cured for at least 24 hours at room temperature prior to imaging.

Primary antibodies used in this study include rabbit anti-tsr (DmCFL, concentration 1:500, gift from Tadashi Uemura (Niwa et al., 2002) against C-terminal peptide CREAVEEKLRATDRQ; rabbit anti-p-Cofilin (concentration 1:500, gift from Tadashi Uemura) against N-terminal peptide (acetyl-A(pS) GVTVSDC; mouse anti-Discs large (concentration 1:100, DSHB 4F3); mouse anti-Bruchpilot (concentration 1:250, DSHB nc82); rabbit anti-GluRIIC (concentration 1:1000, gift from Aaron DiAntonio); mouse anti-GluRIIA (concentration 1:100, DSHB 8B4D2, MH2B); rat anti-Tmod (concentration 1:200, gift from Velia Fowler); guinea pig anti-Coracle (concentration: 1:1500, gift from Richard Fehon).

### Confocal imaging

All samples for comparison were imaged with the same settings between genotypes. Pairs of ventral longitudinal muscles 3 and 4 (muscles 6 and 7) from abdominal hemisegments 2-4 were imaged for all experiments. For *DmCFL* KD, muscle pairs were only imaged if one muscle was class 1 or 2. Z stack images were acquired using an upright Leica Stellaris 5 laser-scanning confocal microscope with dry HC PL Apo 20X/0.75 CS2, oil HC PL Apo 63X/1.40 CS2, or oil HC PL Apo 100X/1.40 CS2 objectives and HyD S detector (Leica Microsystems). Images were acquired sequentially by stack scanning bidirectionally at 400 Hz, with a pixel size of 283.95 nm x 283.95 nm and an area size of 2048 x 1024 pixels in Leica LASX software and saved as LIF files. For all images the pinhole size was 92.53 µm, calculated at 1 A.U. for 561 nm emission. Images were acquired with step sizes of 1 μm (for 20X) or 0.5 μm (for 63X or 100X). For all images for postsynaptic intensity, Z-stack was determined using the HRP channel with the start before and the end 1 μm below the last appreciable HRP signal. FIJI was used to create sum slices or maximum intensity Z projections (NIH).

### Live sample imaging

Live samples were imaged using a Leica Stellaris 5 laser-scanning confocal microscope with a water HC FLUOTAR L VISIR 25X/0.95 objective and HyD S detector (Leica Microsystems). Larvae were dissected and pinned to expose ventral muscles and imaged live in ambient temperature while the larvae were maintained in ice-cold HL3.1 buffer. Images were acquired sequentially by stack scanning bidirectionally at 400 Hz, with a pixel size of 283.95 nm x 283.95 nm and an area size of 2048 x 1024 pixels in Leica LASX software and saved as LIF files. For all images the pinhole size was 92.53 µm, calculated at 1 A.U. for 561 nm emission. Z-stacks were taken using a 1 μm step size and sum slices Z projections were created using FIJI (NIH).

### Structured illumination microscopy (SIM) imaging

All samples were imaged with the same settings on a Zeiss Elyra 7 with Lattice SIM^2^ confocal microscope with a Plan-Apochromat 63X/1.4 Oil DIC M27 objective. Images were acquired sequentially by stack with a pixel size of 0.06 µm x 0.06 µm and an area size of 80.1 µm by 80.1 µm in ZEN Black software and saved as CZI files. Images were taken with 13 phases and reconstructed using ZEN Black software at the “strong” sharpness setting (Zeiss). The Z stacks were acquired at a step size of 0.329 μm and sum slices Z-projections were made using FIJI (NIH).

### NMJ intensity analyses

Postsynaptic intensity measurements were done using Imaris 10.0 (Bitplane). A three-dimensional surface for the HRP source channel was generated for each Z stack image with a surface detail grain level of 0.18 μm, smoothing enabled, and auto-thresholding (Fig. S3A-B). Small surfaces that were not part of the NMJ were removed. For Brp quantification, the sum of the sum intensity of the Brp channel was normalized to the total HRP volume and compared between genotypes. For DmCFL, p-DmCFL, GluRIIC, GluRIIA intensity measurements, an expanded volume was used. A mask was created from the HRP surface using the Distance Transform setting. A second surface was made using the distance transform mask, with a surface detail level of 0.18 and a manual threshold of 0 to 0.5 to limit the surface to a shell extending from the edge of the HRP surface to 0.5 μm away (Fig. S3C-F). The sum of the sum intensity within this expanded surface was normalized to the expanded volume.

### NMJ morphology measurements: bouton counts, NMJ span, subsynaptic reticulum (SSR) area

The Dlg channel was used to quantify NMJ morphology in FIJI (NIH). The number of boutons in a Z-stack image was manually counted using the Cell Counter plugin. NMJ span was measured by drawing a line parallel to the length of the NMJ and normalizing to the length of muscle VL3 (6) or VL4 (7) based on whichever was longer. Cell length was determined by drawing a polygon around the phalloidin-positive shape of each muscle cell and then using the “length” measurement. SSR area was defined as Dlg-positive area normalized to total summed cell areas of muscle 6 and 7. Dlg-positive area was measured by creating a binary mask of the Dlg channel using the Yen thresholding method and recording the area. Cell area was determined by drawing a polygon around the phalloidin-positive shape of each muscle cell and then using the “length” measurement.

### Electron microscopy

Wandering L3 larvae were dissected as described above and fixed overnight in 2.5% glutaraldehyde at 4°C. Larval filets were trimmed to remove the head and tail and to retain the ventral body wall muscles on both sides. Samples were washed three times in 0.1M sodium cacodylate buffer and then post-fixed in 1% osmium tetroxide for 1 hour at room temperature on a rotator. Next, samples were washed three times in 0.1M sodium cacodylate buffer for 10 minutes each wash at room temperature. A dehydration series in ethanol was conducted for 10 minutes at room temperature in each of the following: 30, 50, 70, 85, 95, 100, and 100% ethanol. Infiltration was done in four steps: first in a solution of 1:1 acetonitrile and 100% ethanol for 10 minutes at room temperature followed by infiltration in only acetonitrile for 10 minutes at room temperature. Samples were incubated in a 1:1 mixture of acetonitrile and Embed-812 resin for 30 minutes at room temperature, and then incubated in only Embed-812 resin overnight. In a semi-hardened resin block, larvae were oriented so that the longitudinal edge was along the cutting face of the block. The block was polymerized for 48 hours at 60°C. Thick sagittal sections of 5-10 μm were taken and stained with Toluidine Blue until muscles that were oriented in a longitudinal direction were identified (Fig. S4). A Leica Ultracut UCT ultramicrotome with diamond knife was used to take 70 nm ultrathin sections which were then collected on 3 mm diameter mesh copper grids. Images of individual boutons and the underlying muscle were taken on a JEOL JEM-1400 transmission electron microscope at 100 kV.

### Two-electrode voltage-clamp (TEVC) electrophysiology

TEVC recordings were done on dissected wandering third instars as previously reported (Leahy et al., 2023). Briefly, all recording were done at 18°C in physiological saline (in mM): 128 NaCl, 2 KCl, 4 MgCl2, 1.0 CaCl2, 70 sucrose, and 5 HEPES (pH 7.2). Longitudinally dissected larvae had internal organs removed and peripheral motor nerves cut at the ventral nerve cord (VNC) base. The body walls were glued down (Vetbond, 3M). The preparation was imaged with a 40X water-immersion objective on a Zeiss Axioskop microscope. Ventral longitudinal muscle 6 in abdominal segments 3 or 4 was impaled with two intracellular electrodes (1 mm outer diameter borosilicate capillaries; World Precision Instruments, 1B100F-4) of ∼15 MΩ resistance (3M KCl). The muscle was voltage clamped at −60 mV using an Axoclamp-2B amplifier (Axon Instruments), and the motor nerve stimulated with a fire-polished glass suction electrode using 0.5 ms suprathreshold voltage stimuli at 0.2 Hz from a Grass S88 stimulator. Nerve stimulation–evoked excitatory junction current (EJC) recordings were filtered at 2 kHz. To quantify EJC amplitude, 10 consecutive traces were averaged, and the average peak value recorded. Clampex 9.0 was used for all data acquisition, and Clampfit 10.7 was used for all data analyses (Axon Instruments).

### Statistical analysis

Pairwise comparisons between groups were performed using a two-tailed student’s *t* test with an alpha of 0.05 using R statistical software (R Core Team, 2021). Plots were generated using the ggplot2 R package (Wickham, 2009; Wickham, 2016). Figures report the mean ± standard deviation in addition to sample size.

## Acknowledgements

We thank the Baylies lab members, M. Balakrishnan, M. Lopez, P. Agrawal, A. Beggs and V. Gupta for helpful discussions, and T. Uemura, A. DiAntonio, V. Fowler, and R. Fehon for providing antibodies. The authors thank the Weill Cornell Medicine Microscopy Core Facility and the Bioinformatics Cores at MSKCC for their important contributions to this work. We are grateful to the Bloomington Drosophila Stock Center and the Kyoto Drosophila Stock Center for genetic lines and the Developmental Studies Hybridoma Bank for antibodies.

## Author contributions

Conceptualization: B.C., M.K.B.; Methodology: B.C., S.N.L, D.B.S., V.E.V; Validation: B.C., S.N.L., D.B.S., V.E.V.; Formal analysis: B.C., S.N.L., D.B.S., V.E.V.; Investigation: B.C., S.N.L., D.B.S., V.E.V.; Resources: K.B., M.K.B.; Data curation: B.C., M.K.B.; Writing, original draft: B.C., M.K.B.; Writing & editing: B.C., S.N.L., V.E.V., K.B., M.K.B.; Visualization: B.C.; Supervision: K.B., M.K.B.; Project administration: B.C., M.K.B.; Funding: K.B., M.K.B.

## Data availability

RNA-seq data available using GEO accession number GSE248346.

## Movies

**Movie 1. Individual slices of confocal Z-stack showing *DmCFL* KD Class 2 muscle labeled with phalloidin (red) and HRP (cyan).**

